# Secure Deep Learning on Genomics Data via a Homomorphic Encrypted Residue Activation Network

**DOI:** 10.1101/2023.01.16.524344

**Authors:** Chen Song, Xinghua Shi

## Abstract

Growing applications of deep learning on sensitive genomics and biomedical data introduce challenging privacy and secure problems. Homomorphic encryption (HE) is one of appropriate cryptographic techniques to provide secure machine learning evaluation by directly computing over encrypted data, so that allows the data owner and model owner to outsource processing of sensitive data to an untrusted server without leaking any information about the data. However, most current HE schemes only support limited arithmetic operations, which significantly hinder their applications to support secure deep learning algorithm. Considering the potential performance loss introduced for approximating activation function, in this paper, we develop a novel HE friendly deep network, named Residue Activation Network (ResActNet) to implement precise privacy-preserving machine learning algorithm with a non-approximating activation on HE scheme. We considered a residue activation strategy with a scaled power activation function in the deep network. In particular, a scaled power activation (SPA) function is set within the HE scheme, and so that can be directly deployed on HE computation. Moreover, we proposed a residue activation strategy to constrain the latent space in the training process for alleviating the optimization difficulty. We comprehensively evaluate ResActNet using diverse genomics datasets and widely-used image datasets. Our results demonstrate that ResActNet outperforms other alternative solutions to secure machine learning with HE and achieves low approximation errors in classification and regression tasks.

## Introduction

Security and privacy are growing concerns in various domains of data science as well as in general machine learning (ML). These concerns arise from both data and model perspectives. On the one hand, data themselves (e.g., financial or biomedical data) can be sensitive or susceptible to privacy leakage or security attacks. On the other hand, ML models or data analytical tools (especially those based on modern deep and complex models) can be vulnerable to adversarial attacks, which put the model parameters functions, and privacy of training samples at risk Ribeiro et al. [2015], De Cristofaro [2020], Shokri et al. [2017], Fredrikson et al. [2015, 2014], Tramèr et al. [2016], Ateniese et al. [2015], Shringarpure and Bustamante [2015]. In real-world scenarios, the privacy and security issues of data and models can be intertwined and make it challenging to develop or release secure and privacy-preserving ML models on data of various types including sensitive or even confidential data. Typically, there are four widely-used approaches to privacy-preserving machine learning. First, multi-party computation allows multiple parties to jointly evaluate functions while concealing information from outside parties De Cristofaro [2020]. Second, another strategy for privacy protection and preservation is differential privacy (DP), which aims to guarantee an algorithm to learn the statistical information of the population without disclosing information about individuals via adding noises to data statistics Choudhury et al. [2019], Kim et al. [2019], Cormode et al. [2018], Chen et al. [2020], Tramèr et al. [2015]. Third, a cryptographically-protected hardware, such as Intel SGX McKeen et al. [2013], protects private data by executing codes in a trusted execution environment Rushby [1981]. Alternatively, homomorphic encryption (HE) technique provides a solution by allowing computation directly over encrypted data when sensitive data needs to be evaluated by untrusted parties. In this research, we focus on the homomorphic encryption techniques.

A homomorphic encryption (HE) scheme directly performs computation on encrypted data to obtain encrypted results. After decryption on the results, the decrypted results can match the outcome from the corresponding operations on the plaintext. However, one of the notorious drawbacks of existing HE schemes is that they support a rather limited set of operations (e.g. homomorphic additions and multiplications), and can not implement other common operations like comparisons and exponential operations. Hence, these HE schemes cannot support deep and complex models with satisfying performance and HE requirement. For example, rectified linear units (ReLU) and Sigmoid functions, commonly used as nonlinear activation functions in deep learning (DL) models, cannot be computed using simple operations provided by existing HE schemes within HE’s polynomial requirement. To deal with this constraint, polynomial approximation is widely employed as the complex activation functions in practice. Previous works Ishiyama et al. [2020], Hong et al. [2021], Chen et al. [2018], Al Badawi et al. [2018] employed high-degree approximation of ReLU and Sigmoid function to deep learning to improve image classification on MNIST and CIFAR10 datasets. However, there are two major limitations in utilizing polynomial approximation of ReLU and Sigmoid functions. First, recent approximation method can only accurately approximate ReLU and Sigmoid function within a limited range, and thus batch normalization must be used in polynomial approximation of DL model, which increases computation cost. Second, although polynomial approximation works well on classifications, the inevitable approximation error introduced significantly affects the performance of regression task. Other approaches to implement activation in DL model include using a square activation without approximation Al Badawi et al. [2018], Benaissa et al. [2021], Chabanne et al. [2017], Hesamifard et al. [2017]. However, square activation does harm to classification performance and has an optimization problem in training stability compared with polynomial approximation.

In this paper, we develop a novel HE scheme to address the nonlinear mapping issues in deploying secure deep models utilizing HE. Instead of trying to reduce the approximating error to estimate the HE-unfriendly activation function from polynomial approximation, we explicitly designed the plain model within HE friendly operations. More specifically, we built a residue activation layer to fit the nonlinear mapping in hidden layers of deep models. In detail, by employing scaled power activation (SPA) function merely on the part over the global average value, we assumed that the residue activation layer can better fit the nonlinear mapping robustly in a HE-friendly manner, compared with the original square activation layer. We thus built a deep network, named ResActNet (Residue-activation Network), with ResActNet activation layers. In the ResActNet, we employed a scaled power function as nonlinear activation, where a scalar term is worked for tuning the convergence of network. To tackle the unbounded derivative issue of the scaled power function, we deployed a residue activation strategy in the network. In detail, we constraint the SPA on the residue of latent vector in deep network. In this way the divergence of network can be avoided. It is noticed that the SPA and residue activation strategy can be conducted within addition and multiplication, so that it can be directly deployed to a fully HE scheme without approximation. In this regard, the approximating error is avoided theoretically. Experimental results on multiple datasets show that ResActNet works well on different prediction tasks including classification and regression.

To sum up, innovative contributions of this study can be viewed from the following three perspectives.

**Firstly**, we reduced the performance loss of HE evaluation via deploying a non-approximating HE scheme. By setting a novel nonlinear function named scaled power activation, a deep network in our proposed ResActNet is able to implement HE without approximating nonlinear activation functions. Meanwhile, unlike the polynomial approximation method that has a specific effective approximation range requirement for the input, the deep model with proposed non-approximating scheme in ResActNet can accurately evaluate deep models without any range limitation. From this perspective, the proposed ResActNet method has excellent generalization performance compared with alternative approximating methods.

**Secondly**, we designed a novel network structure, named the residue activation layer, to solve the optimization problem of power activation function in HE evaluation. Instead of directly performing nonlinear mapping in the hidden layers using a square function, which has been proven to be unstable and hardly optimized, we only add nonlinearity to a residue part of hidden tensor. By doing this, the divergence problem in a HE-based deep model can be alleviated by constraint the affects of power activation function.

**Finally**, we set a more flexible and powerful scaled power activation (SPA) function for our proposed HE scheme. Experimental results show that tuning the scalar term and power order in a deep model can improve the model’s performance.

## Methods

In this section, we describe the approach we specifically designed to improve secure and privacy preserving evaluation of machine learning models with HE scheme. More specifically, we design a flexible HE friendly activation function for nonlinearly mapping hidden feature in the deep model, and implement a special constrainted activation strategy (i.e., residue activation layer) to achieve stable optimization with the scaled power activation (SPA) function. Combining with widely used CNN and dense layer, we developed a new network, partial activation network (ResActNet), and showed that it can be directly deployed to an HE scheme for providing secure machine learning on various tasks and models.

As shown in **Fig.1,** we assume that information leakage may happen when the model evaluates the data on the untrusted party (e.g., a commercial online server). Aiming to avoid information leakage, a model owner may encrypt model parameters *m* as [*m*], and data owner typically encrypt data *x* as [*x*], before sensitive information being transferring to untrusted party. So, the evaluation between [*m*] and [*x*] operated in encrypted form enabled by HE technique, and return an encrypted result [*r*]. Before the [*r*] being transferred to data owner and then decrypted, nobody can access to the result in a plain form, so that the information privacy is protected.

**Fig. 1.**
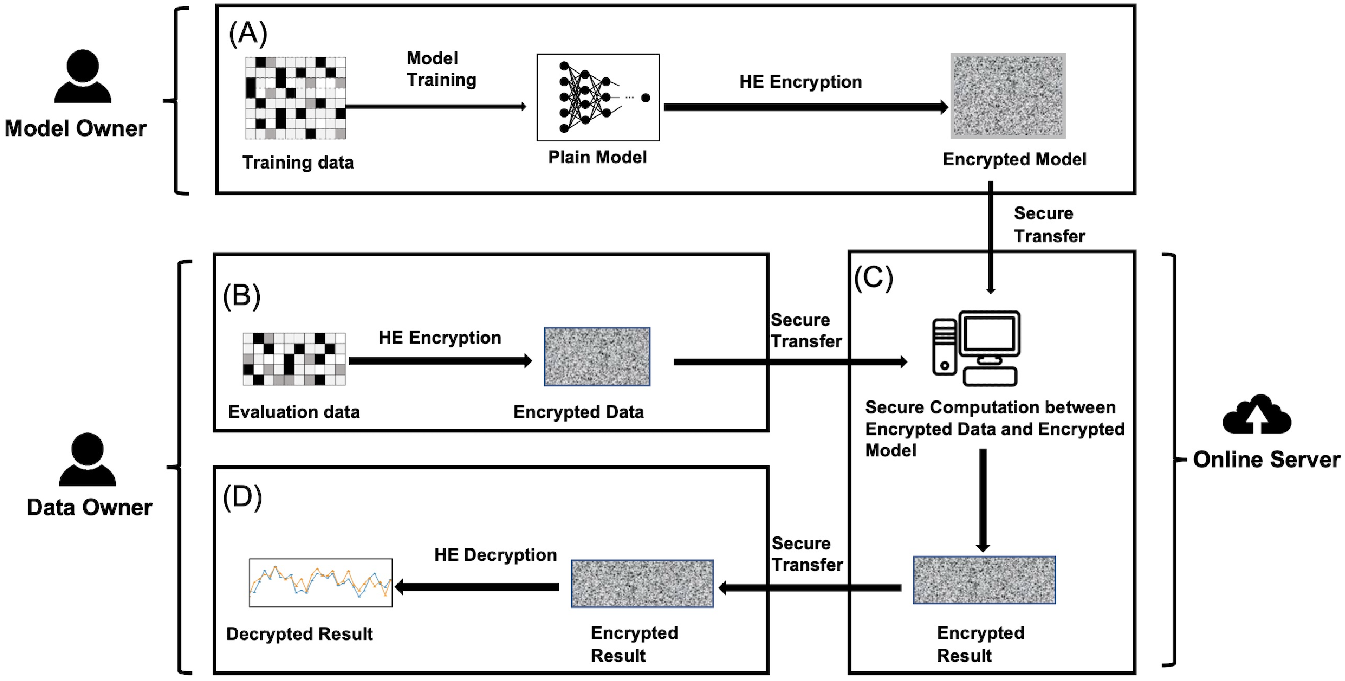
Overview of our privacy-preserving ML methods. (A) Model owners train a plaintext model in the HE scheme and encrypt the model parameters. (B) Data owner encrypts data before a piece of data is transferred for being evaluated by an untrust third party. (C) The machine learning computations perform directly on encrypted data and return an encrypted results. (D) The returned encrypted results can only be decrypted by data owner. The private data will never be exposed to anyone in unencrypted form so that the privacy is preserved.

The overall structure of ResActNet is depicted in **Fig.2.** We utilized a widely used CNN layer Benaissa et al. [2021], Obla et al. [2020a], Ishiyama et al. [2020] for extracting latent information of features. After the CNN layer, we proposed a partial activation layer to map hidden features to a nonlinear space. In the residue activation layer, a HE friendly operation, SPA function is deployed on the residue part of hidden tensor. It is noticed that we constructed the residue activation layer with merely additions and multiplications. In this perspective, ResActNet can be directly implemented with HE.

**Fig. 2.**
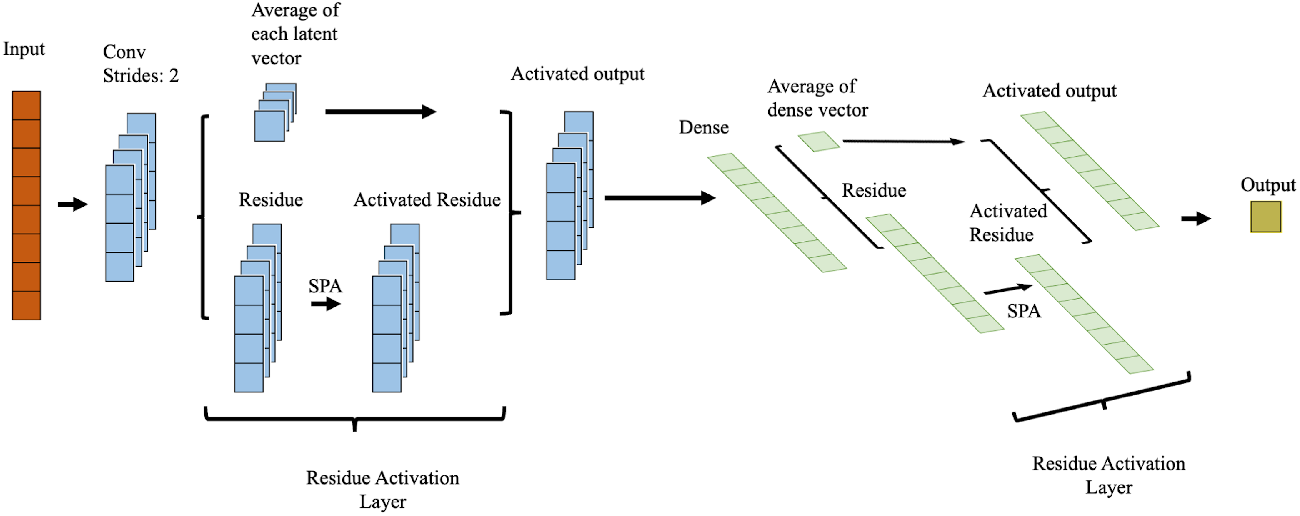
Model structure of the proposed ResActNet for HE evaluation. In ResActNet, we map hidden features non linearly using a novel residue activation layer, after the CNN layer or dense layer. In this residue activation layer, the scaled power activation (SPA) function is used to add non-linearity to the residues of hidden features to reduce divergence resulted from the unbounded derivative of the SPA function. Since the residue activation layer is constructed with merely additive and multiplicative operations, it is feasible to directly employ fully HE schemes on ResActNet and thus no approximation error is included during HE.

### Scaled Power Activation

The primary role of the activation layer is to capture the non-linearity of complex features in a deep model. Complex tasks such as classifying images, genomics or speech involve the separation of non-linear data and can be conducted using non-linear models. For example, non-linear activation functions including Rectified Linear Unit (ReLU) and Sigmoid enable neural networks to perform this type of task. Nonetheless, those well-performed non-linear activation functions fail to satisfy the requirement of fully HE scheme in existing work, where only additions and multiplications are homomorphically supported. Therefore, in this study, we propose a scaled power activation function to nonlinearly map hidden layers in deep models.

Denoting the input of an activation function as *z*, an activation function is represented as *σ*(*z*). The scaled power activation function can be formalized as

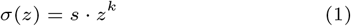

where *s* is the scalar term and *k* is the power term which adds nonlinearity to hidden features. This scaled power activation function satisfies the continuous and differentiable qualities of activation. Noting that the scaled power activation is not necessarily monotonic which makes it hard to train Parascandolo et al. [2016]. So we set a scaled term for scaling hidden features so that they are zero-centered and can be effectively optimized Le Cun et al. [1991].

It is noticed that another potential issue of the SPA function is the unbounded derivative. Easily, we have the derivative of Eq.(1) is 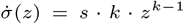. Existing works showed that it is hard to optimize a model that has activation function with unbounded derivative. To solve this problem, we designed a partial activation method to alleviate the optimization problem of SPA function, which is describe in detail in the next section.

### Residue Activation Layer

Although using a scaled power activation function ensures that an efficient deep model can be implemented with non-linear activation functions, it is still challenging to efficiently optimize a deep network equipped with a scaled power activation function since the unbound derivative of SPA. Inspired by the residue network Targ et al. [2016] and a truncation trick in one-shot learning generative network Yang et al. [2021] to handle the divergence problem in deep neural networks, we propose a partially activation strategy (i.e., residue activation layer) for deep networks as follows. More specifically, we only activate the part of hidden features that are greater than a global average value. Since the global average value remains constant, it is easy to reduce divergence resulted from unbounded derivatives.

Mathematically denoting a layer as *L*, and the input tensor of one layer as *z*, the partially activation layer can be written as

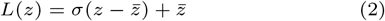

in which *σ*(·) is the scaled power activation function and 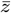 is the average value of hidden tensors. In Eq.(2), hidden features *z* have been split into the global average part 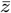 and the residue part 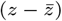. Only the residue part is added non-linearity from the SPA function so that the unbounded derivative issue only effects the residue part. Besides, the gradient in corresponding hidden layers would remain small and is slightly updated. In this respect, the model equipped with partial activation layer is more stable and robust.

Note that all operations in the partially activation layer can be implemented via multiplications and additions. In this way, the partially activation layer can be directly transferred to an HE scheme for secure evaluation without any approximation, and thus the consistence of model performance is ensured.

#### Algorithm 1 Optimizing a residue activation layer with the scaled power activation function in ResActNet

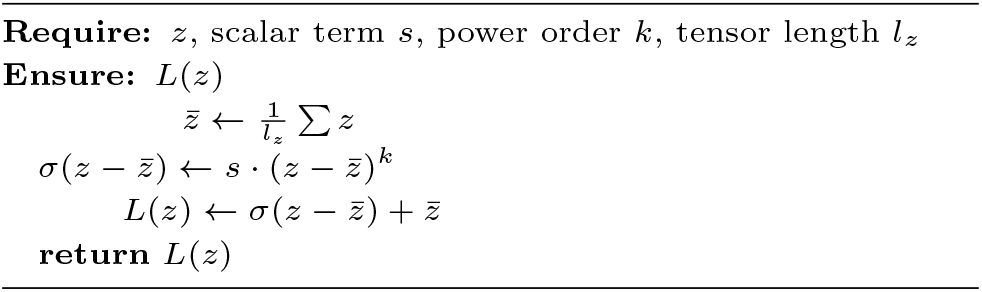

### Non-approximating HE Evaluation via ResActNet

We built ResActNet based on a scaled power activation function and a residue activation layer. In detail, We combined the partial activation layer with a CNN layer and a dense layer, which have already been proved to be HE-friendly and have been widely used in HE evaluation Benaissa et al. [2021], Obla et al. [2020a,b]. ResActNet consists of a stack of CNN blocks for capturing hidden features and a partial activation block for nonlinear mapping as well as global pooling. To implement a convolution layer in ResActNet, we set the strides as 2 in a plain model to downsample the feature space as most HE-based CNN models used Obla et al. [2020a,b]. We set two hyperparameters in the residue activation layer, which are a scalar term and a power order term to control the convergence of a model. **Fig.2** shows an example of ResActNet, where residue activation layers follow with CNN layers for nonlinear mapping. Compared with the infrastructure of approximation network in Panel II of **Fig.1,** the network in ResActNet (**Fig.2)** is constructed with merely additions and multiplications, so that it can be directly implemented in an HE scheme without any approximation.

## Experimental Results

In order to demonstrate the effectiveness, we conducted comprehensive experiments to evaluate our proposed method on multiple datasets including a quantitative genomics dataset for evaluating the regression performance, a TCGA dataset for multilabel tumor type prediction and a Sars-CoV-2 dataset for strains classification task. We compared the proposed method with varies baseline methods, including linear regression, support vector machine, multi-layer perceptron, to show the performance of the proposed model.

### Datasets

We chose genomic datasets not only because genomic data usually contains sensitive information susceptible to privacy leakage, but also because genomic data is typically at large scale and with high dimension (i.e. features significantly outnumber samples). In addition to having high volume and sparsity, genomic data is also complicated in that features are typically highly correlated and there are interactions and complex relationships among them. We selected one human genomic dataset from the Cancer Genome Atlas (TCGA) dataset Tomczak et al. [2015] to predict cancer subtypes from gene expression profiles. We picked another distinct genomic dataset from the Severe acute respiratory syndrome coronavirus 2 (SARS-CoV-2) sequencing Wu et al. [2020] to classify virus strains based on their sequences. Since all the other datasets have categorical labels, we selected a widely-used yeast genomic dataset Bloom et al. [2015] with quantitative phenotypes to evaluate the performance of our proposed approach on a regression task.

#### Genomics data

##### TCGA

Our experiments are conducted on a sub-dataset from The Cancer Genome Atlas (TCGA) dataset. We collected somatic mutation data from 21,199 points and copy number variants data from 25,128 gene points of 2,713 samples with 11 tumor types. We used the occurrence of a gene point as the feature value, in which 1-value represent somatic mutation found in a gene point and 0-value represent no somatic mutation founded in a gene point. In copy number variant part, we labeled deletion as −1 and −2, and we labeled amplification of a gene as 1 and 2 value in a copy number variants, 0 means no copy number variants founded yet. We concatenated two parts of mutation into a feature matrix for predicting 11 tumor positions. An *L*1 normalized logistic regression model was employed to selected 1000 features from the raw data with penalty equals to 0.1. **SARS-CoV-2:** SARS-CoV-2 causes the world-wide epidemic and have mutated rapidly. It is essential to sequence the viral sample from a patient and classify it into one of the known strains for managing the epidemic. We used four strains of genomes data, which is ‘B.1.427’ (Epsilon), ‘B.1.1.7’ (Alpha), ‘P.1’ (Gamma), ‘B.1.526’ (Iota). Each strain contains 2000 samples. We encoded the genomes data {*N*, *A*, *G*, *C*, *T*, *K*, *W*, *Y*, *R*, *M*, *S*} as {0, 1, 2, 3, 4, 5, 6, 7, 8, 9, 10}. We selected 500 features via an L1 normalized LR model with the penalty term equal to 10. **Yeast:** The yeast dataset Bloom et al. [2015] contains categorical genotypes of 28,820 unique genetic variants that were obtained by sequencing 4,390 individuals from a cross between two strains of yeast: a widely used laboratory strain (BY) and an isolate from a vineyard (RM). We encoded the features as 0 for genotypes from BY and 1 for those from RM, and normalized the phenotypes to be used as quantitative labels. LASSO can be used to select features since it can constrain the sparsity of weight matrix, and corresponding features with the non-zero weights decide the decision hyperplane, so that they can be treat as important feature Fonti and Belitser [2017]. A LASSO model with parameter is used to select features, using *α* = 0.001 learned from cross-validation (Appendix Table.7).

### HE-based secure prediction

We investigated the proposed method for different prediction tasks on multiple dataset. Three models were employed here as baseline, namely, Linear regression(LR), support vector machine (SVM) and multi-layer perceptron (MLP), respectively. We built the LR and SVM model with the scikit-learn package Pedregosa et al. [2011] while building the MLP model and the proposed model with TensorFlow Abadi et al. [2015] on a NVIDIA Tesla Volta V100 GPUs with 16GB RAM. We used a quantitative accuracy as a metric to learn hyperparameters of the model for classification model. We used the mean square error (MSE) as a quantitative metric to learn hyperparamters of regression models. As a measurement for selecting optimal hyperparamters of different model on multiple datasets, the average accuracy or MSE from crossvalidation of each hyperparameter were used. We implemented HE using the Cheon-Kim-Kim-Song (CKKS) scheme. In CKKS, homomorphic addictions and multiplications are operated with a vector form with real values. Firstly introduced in Cheon et al. [2017], CKKS is a Ring learn-with-error (RLWE) approximate arithmetic to employ HE operations. Since CKKS can compute element-wise homomorphic multiplication and addition over encrypted vectors within predeterminated precision, it is widely used in secure and privacy preserving ML. We implemented the CNN inference program over HE using the Tenseal package Benaissa et al. [2021], which is implemented with Microsoft’s Simple Encrypted Arithmetic Library (SEAL) SEAL on the CKKS scheme with a security level greater than 128-bit Albrecht et al. [2021]. And the HE encrypted computations were conducted on 40 threads of Intel Xeon (Cascade Lake) CPU cores with 1.5TB of RAM in total.

**Table.1** summarizes the HE prediction accuracy of ResActNet compared with competing methods. We observed that the ResActNet outperform the other widely used homomorphic encryption machine learning models in a view of classification accuracy and regression mean square error.

**Table 1.** Prediction results on multiple datasets for classification and regression. **LR**: linear regression; **SVM**: support vector machine; **MLP**:multi-layer perceptron; **ResActNet**:the proposed residue activation Network. For classification task, the 10 repeats of average accuracy (larger is better) for 10-label tumor classification (TCGA) and 4-strain virus classification are listed in the table while for regression task, the 10 repeasts of average mean square error (MSE, smaller is better) for predicting two quantitative yeast phenotype from genotype are listed in this table.

| Method | Classification |  | Regression |  |
| --- | --- | --- | --- | --- |
|  | Covid | TCGA | Cobalt | Copper |
| **LR** | 0.9855 | 0.8286 | 0.0118 | 0.0074 |
| **SVM** | 0.9875 | 0.8283 | 0.0115 | 0.0175 |
| **MLP** | 0.9775 | 0.7574 | 0.0118 | 0.0117 |
| **ResActNet** | **0.9887** | **0.8382** | **0.0086** | **0.0041** |

### Scalar term in SPA for controlling convergence

The scalar term set in SPA is for controlling the convergence of loss in a deep network. We investigated the effects of different scalars on the training and validation datasets to predict labels in TCGA, in terms of cross-entropy loss between the training and test dataset. **Fig.3** shows effects of different values of scalar in the power activation. Divergence occurred when a normal square activation function was used, in which no scalar term was applied. Oscillation was alleviated with decreasing the value of scalars. However, if we decreased the scalar to 1*e*^−4^, the loss function would be hard to converge. This results indicated that the scalar term can be deployed in HE model for tuning the convergence.

**Fig. 3.**
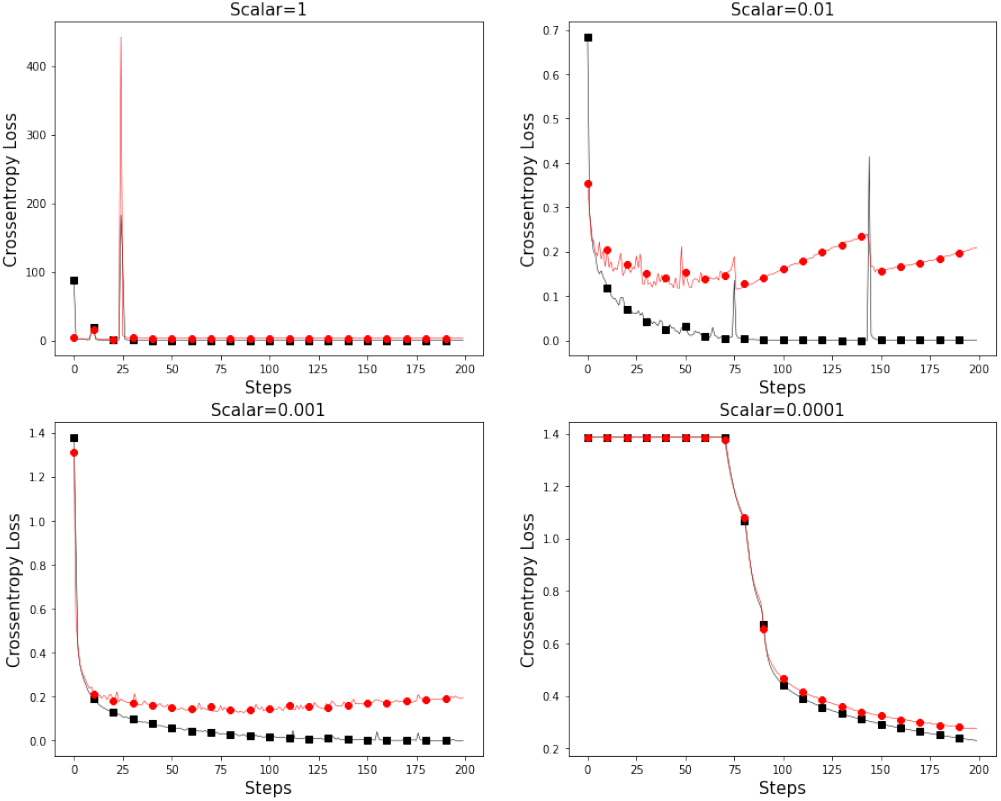
The effects of scalar in the scaled power activation function. The crossentropy loss in 200 training steps of different scalar are shown. The red line is the crossentropy loss of evaluation dataset, while the black line is that of corresponding training dataset.

### Evaluation of Homomorphic Encryption on ResActNet

In this section, we evaluated the performence loss introduced from homomorphic encryption technique with difference parameters for secure machine learning on genomics data. In detail, we evaluated the homomorphic encryption method in two perspects, **1)** the gross computation cost of HE machine learning; **2)** the difference between decrypted output from a HE model with output from corresponding plain model (HE evaluation). For the classification task, we used accuracy between the decrypted output with the plain results as a metric to evaluate the HE method, while for the regression task, we used the mean square error and correlation score as a measurement to evaluate the performance of HE method. **Table. 2** shows the model evaluation with corresponding parameters in CKKS HE scheme. The performance of HE regression on the yeast data is in **Table.2.(a)** and the HE evaluation on classification on TCGA and Covid datasets are in **Table.2.(b)** and **Table.2.(c)** *N* and *q* are the polynomial degree and *q* the ciphertext modulus parameters,respectively and determine the security level and HE precision. The value of *logq* determines the times of homomorphic multiplication on the ciphertext, the larger *logq* is, the more times of multiplication the HE can operate. We set the *logq* and *N* according to the HE standard Albrecht et al. [2021]. It is shown that there is a tradeoff between the security level, performance and computation cost. The time burden increased when we encoded the data into a large polynomial modulus (i.e., a larger value of *N*). Using a larger value of bit scale can improve the precision of HE scheme, while harming the security level.

**Table 2.**
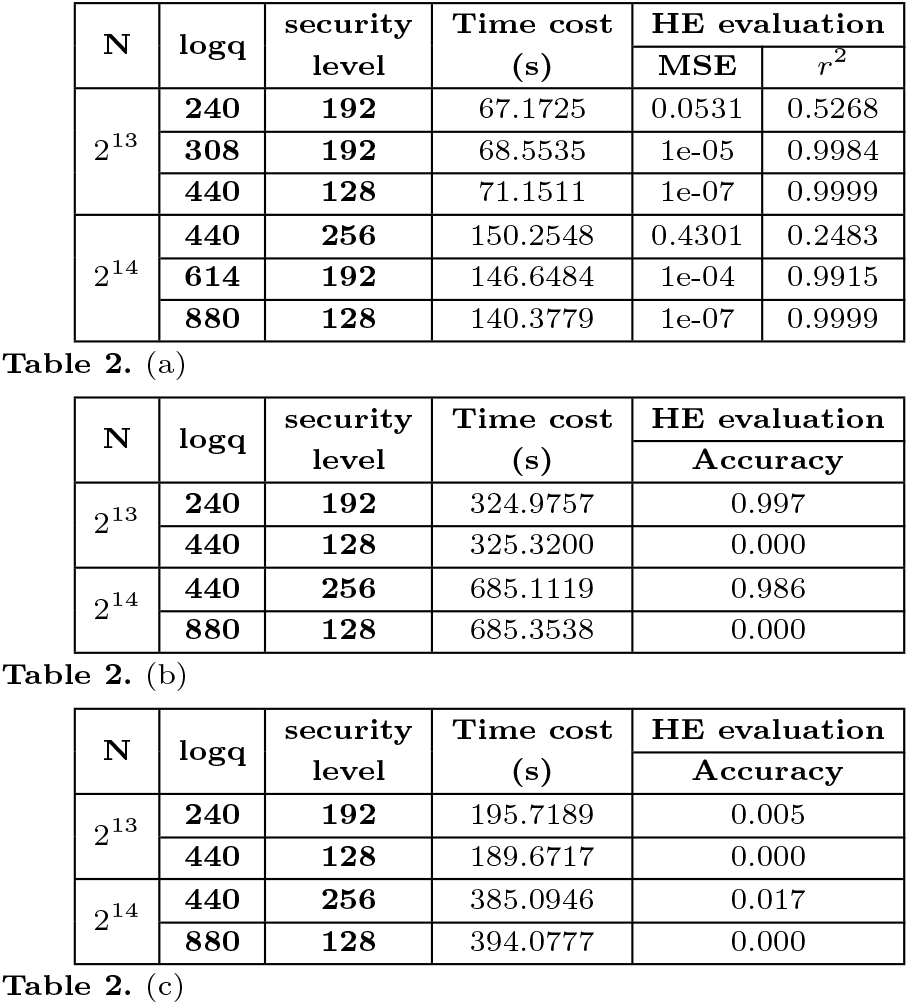
Comparison of different homomorphic encryption parameters. *N* is the polynomial degree of CKKS scheme; and logq is a parameter used in each homomorphic multiplication; Time cost is the total time burden for encryption, computation over encrypted data, and decryption per 100 samples. (a) HE evaluation of yeast phenotype regression task. The mean square error (MSE) and *r*^2^ score between results from a plain-text model and the HE model on the same input data are listed to evaluate the HE scheme.(b) and (c) TCGA multilabel tumor classification task and Sars-cov-2 (COVID) strain classification task, respectively. The accuracy between results from a plain-text model and the HE model on the same input data are listed to evaluate the HE scheme.

## Conclusion

In this study, we improved secure deep model evaluation via HE by setting a non-approximating scheme named ResActNet. Since existing fully HE schemes only support homomorphic addition and homomorphic multiplication, we employed a scaled power activation in hidden layer to add nonlinearity. In order to solve the divergence problem caused by the unbounded derivative of scaled power activation, we proposed a partial activation approach to constrait the scaled power activation on residue of hidden features. We deployed the partial activation method with a CNN block and dense block and proposed a ResActNet for HE evaluation of classification and regression tasks. We evaluated the proposed ResActNet on multiple tasks, including multi-label tumor type prediction using TCGA data, Sars-Cov-2 virus strain classfication task, and prediction of quantitative traits using a yeast genomic dataset. Our results show that the deep model with partial activation strategy in ResActNet outperforms baseline and alternative methods. Moreover, since ResActNet can be directly implemented with HE schemes, it outperforms alternative methods without introducing approximation errors when implementing HE.

Despite the success of the proposed method, there are several interesting directions to explore to further improve these strategies. For example, we will explore HE-based training strategies which is hampered by the model complexity. Additionally, more advanced HE algorithms should be explored to mitigate potential risks of information leakage caused by intermediate model parameters exchanged in federated learning or distributed learning settings Sav et al. [2022], Nasr et al. [2019], Hitaj et al. [2017], Wang et al. [2019].

In addition to technical approaches that have been discussed in this paper, there are regulations and laws established to protect data privacy including the Health Insurance Portability and Accountability Act (HIPAA) Assistance [2003] in the US and the EU General Data Protection Regulation (GDPR) Voigt and Von dem Bussche [2017] in Europe. However, these existing laws and regulations significantly lag behind technical advances and do not fully protect data generated from third parties. We believe that the proposed methods in this study point to a future direction in developing efficient secure ML algorithms deployed on massive datasets in a variety of data science domains.

## Acknowledgements

We would like to acknowledge the organizers of IDASH PRIVACY & SECURITY competition for providing TCGA subset dataset and Sars-CoV-2 sequencing dataset in 2020 iDASH and 2021 iDASH competitions, respectively.

